# In vitro discovery and optimization of a human monoclonal antibody that neutralizes neurotoxicity and lethality of cobra snake venom

**DOI:** 10.1101/2021.09.07.459075

**Authors:** Line Ledsgaard, Andreas H. Laustsen, Urska Pus, Jack Wade, Pedro Villar, Kim Boddum, Peter Slavny, Edward W. Masters, Ana S. Arias, Saioa Oscoz, Daniel T. Griffiths, Alice M. Luther, Majken Lindholm, Rachael A. Leah, Marie Sofie Møller, Hanif Ali, John McCafferty, Bruno Lomonte, José M. Gutiérrez, Aneesh Karatt-Vellatt

**Affiliations:** Department of Biotechnology and Biotherapeutics, Technical University of Denmark; DK-2800 Kongens Lyngby, Denmark; IONTAS Ltd.; Cambridgeshire CB22 3FT, United Kingdom; Sophion Bioscience, DK-2750 Ballerup, Denmark; Instituto Clodomiro Picado, Facultad de Microbiología, Universidad de Costa Rica; San José 11501-2060, Costa Rica; Quadrucept Bio, Cambourne CB23 6DW, United Kingdom

## Abstract

The monocled cobra (*Naja kaouthia*) is one of the most feared snakes in Southeast Asia. It is a highly dangerous species with a potent venom deriving its toxicity predominantly from abundant long-chain α-neurotoxins. The only specific treatment for snakebite envenoming is antivenom, which is based on animal-derived polyclonal antibodies. Despite the lifesaving importance of these medicines over the past 120 years, and their ongoing role in combating snakebite disease, major limitations in safety, supply consistency, and efficacy creates a need for a new generation of improved treatments based on modern biotechnological techniques. Here, we describe the initial discovery and subsequent optimization of a recombinant human monoclonal immunoglobin G (IgG) antibody against α-cobratoxin using phage display technology. Affinity maturation of the parental antibody by light chain-shuffling resulted in an 8-fold increase in affinity, translating to a significant increase in *in vitro* neutralization potency and *in vivo* efficacy. While the parental antibody prolonged survival of mice challenged with purified α-cobratoxin, the optimized antibody prevented lethality when incubated with *N. kaouthia* whole venom prior to intravenous injection. This study is the first to demonstrate neutralization of whole snake venom by a single recombinant monoclonal antibody. Importantly, this suggests that for venoms whose toxicity relies on a single predominant toxin group, such as that of *N. kaouthia*, as little as one monoclonal antibody may be sufficient to prevent lethality, thus providing a tantalizing prospect of bringing recombinant antivenoms based on human monoclonal or oligoclonal antibodies to the clinic.

**One Sentence Summary:** A recombinant human monoclonal immunoglobulin G antibody, discovered and optimized using *in vitro* methods, was demonstrated to neutralize the lethal effect of whole venom from the monocled cobra in mice via abrogation of α-neurotoxin-mediated neurotoxicity.

## INTRODUCTION

Snakebite envenoming is a neglected tropical disease, which each year claims hundreds of thousands of victims, who are either left permanently disfigured or meet an untimely death *(1)*. Asia is the continent where most bites and deaths occur. In Southern and Southeast Asia, the monocled cobra (*Naja kaouthia*) is responsible for a large number of the recorded severe snakebite cases *(2, 3)*, which is exemplified by the fact that 34% of snakebite-related deaths in Bangladesh from 1988 to 1989 were attributed to bites from either *N. kaouthia* or the closely related species, *N. naja (4)*. Life-threatening clinical manifestations of *N. kaouthia* envenomation include flaccid paralysis due to the actions of abundant long-chain α-neurotoxins, which block neuromuscular transmission by binding to the nicotinic acetylcholine receptor (nAChR) with high affinity, causing a curare-mimetic effect *(5, 6)*. These long-chain α-neurotoxins belong to the three-finger toxin superfamily, which dominate the venom in terms of abundance and toxicity, as judged by their high Toxicity Scores *(6–8)*, and are thus the main toxin targets to be neutralized for successful intervention in human snakebite cases.

Each year, the lack of access to affordable and effective treatment against snakebite envenoming leaves thousands of victims in despair. Animal-derived antivenoms remain the cornerstone of snakebite envenoming therapy *(1)* and are still produced by immunizing large mammals, usually horses, with snake venom, followed by the purification of antibodies from the blood plasma, resulting in polyclonal antibody preparations *(9)*. Being heterologous products, animal-derived antivenoms often lead to a range of adverse reactions whose incidence varies depending on the product *(10)*. Furthermore, it is estimated that only a fraction of the antibodies in current antivenoms contribute to neutralization of relevant toxins. Large amounts of antivenom are therefore required to treat a snakebite case, resulting in heterologous protein loads as high as 15 grams per treatment in severe envenoming cases *(11, 12)*. Moreover, antivenoms targeting elapid species in particular have a low content of therapeutic antibodies against low molecular weight neurotoxins with high Toxicity Scores, as a discrepancy exists between the toxicity (high) and the immunogenicity (low) of these toxins, leading to weaker immune responses in production animals *(13–16)*. Despite many advances within antibody technology and biotechnology, a need remains for antivenoms with improved safety and efficacy *(17, 18)*.

Recently, recombinant antivenoms based on oligoclonal mixtures of human antibodies have been proposed as a cost-competitive alternative to current antivenoms *(19–22)*. Such recombinant antivenoms may offer safer and more efficacious snakebite envenoming therapy due to their compatibility with the human immune system and the possibility of only including efficacious antibodies, targeting medically relevant snake toxins, in the antivenom mixture *(18)*. Moreover, it has been demonstrated that such oligoclonal recombinant antivenoms, consisting of carefully selected immunoglobulin Gs (IgGs), can be developed using phage display technology *(19)*. However, to date, no neutralizing human monoclonal IgG has been reported against α-neurotoxins, which are among the medically most important toxins present in elapid venoms. Additionally, no recombinant antibody has yet been reported to neutralize whole venom from any animal following intravenous injection.

Here, we report the discovery and affinity maturation of a human monoclonal IgG targeting α-cobratoxin, the medically most relevant toxin from *N. kaouthia* venom. Using chain-shuffling, the affinity of the antibody was improved 8-fold, resulting in enhanced neutralization both *in vitro* and *in vivo*. The increased affinity to α-cobratoxin improved the IgG from one that was able to prolong survival of mice challenged with purified α-cobratoxin to one that could neutralize the lethal effect of whole *N. kaouthia* venom in mice.

## RESULTS

### A phage display derived fully human antibody that prolongs survival *in vivo*

A naïve scFv phage display library *(23)* was used to carry out three rounds of panning against biotinylated α-cobratoxin from *N. kaouthia* immobilized on a streptavidin coated surface. Antibody-encoding genes (scFv format) were isolated from both the second and third panning rounds and subcloned into a bacterial expression plasmid *(24)*. In total, 282 clones harboring this expression plasmid were picked for antibody expression and subjected to binding analysis by DELFIA as previously described *(19)*. Of these, 36 clones displaying specific binding signal against α-cobratoxin (25 times above the background binding signal) were picked for DNA sequencing and further characterization (Fig. 1A).

**Fig. 1.**
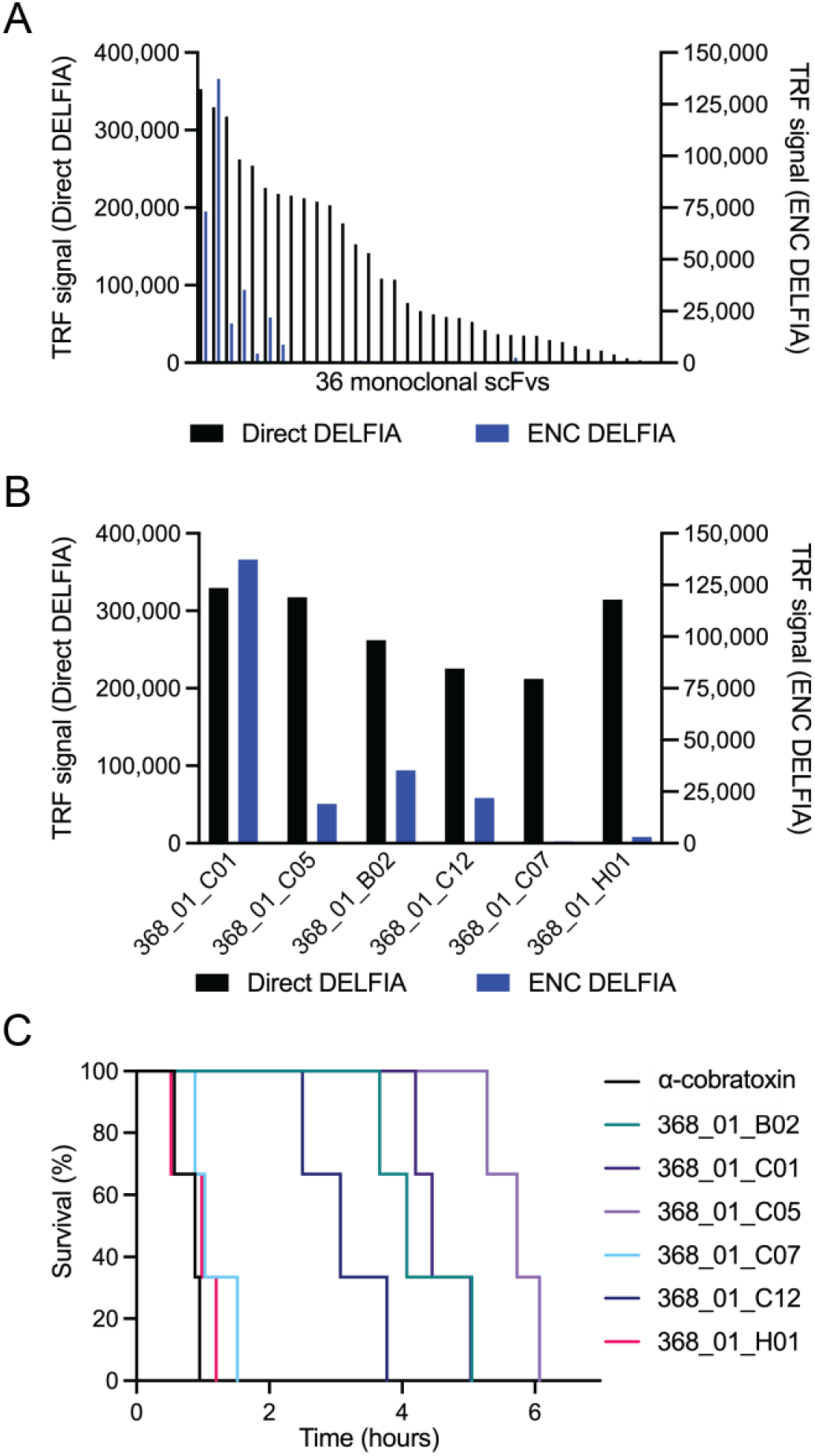
Affinity ranking of scFvs and Kaplan-Meier survival curves for mice co-administered with IgG antibodies and α-cobratoxin. (**A**) Direct and ENC DELFIA of 36 monoclonal scFv-containing supernatants. (**B**) Direct and ENC DELFIA of the top six monoclonal scFv-containing supernatants. (**C**) Kaplan-Meier survival curves for mice co-administered with IgG antibodies and α-cobratoxin. α-cobratoxin refers to mice injected with α-cobratoxin alone.

A total of 28 scFvs with unique V_H_ and V_L_ CDR3 regions were evaluated in an expression-normalized capture (ENC) assay, which eliminates the expression variation between the clones and allow ranking based on affinity. The six α-cobratoxin-binding scFvs that yielded the highest binding signals (Fig. 1B) were selected for reformatting into the IgG format and transiently expressed using Expi293F™ cells. All six α-cobratoxin-targeting antibodies retained binding to α-cobratoxin upon conversion. However, when the antibodies and α-cobratoxin were preincubated and administered i.v. to mice, the antibodies failed to prevent lethality, although they did succeed in prolonging survival significantly (Fig. 1C). One plausible explanation for the limited efficacy of these antibodies could be due to their sub-optimal binding affinity to α-cobratoxin.

### Affinity maturation using chain-shuffling

*In vitro* affinity maturation strategies involve two key steps, diversification of the primary antibody sequence and enrichment of affinity-improved antibody variants using a selection platform such as phage display technology. Diversification of primary antibody sequence can be achieved by introducing mutations to the variable regions using random or targeted mutagenesis. Alternatively, new combinations of heavy and light variable regions can be made by recombining selected heavy or light chains with a repertoire of partner chains by a process known as chain shuffling *(25)*. Here, the 368_01_C01 antibody was chosen for further study due to its combined performance in the ENC DEFLIA and *in vivo* experiments. This antibody was therefore subjected to light chain shuffling to diversify sequence by pairing the V_H_ chain with a library of naïve kappa and lambda light chains. Following library generation, three rounds of stringent panning against α-cobratoxin were completed. For precise control of antigen concentration, phage display selections were carried out in solution phase. The phage antibodies were allowed to bind to biotinylated α-cobratoxin in solution, and the bound phage was subsequently captured using streptavidin-coated beads for washing and elution. The antigen concentration was lowered 10-50-fold in each round to selectively enrich antibodies with high affinity. The polyclonal phage outputs for the selections were tested for binding to α-cobratoxin through polyclonal DELFIA, revealing that α-cobratoxin-binding scFvs were present in all three rounds, while negligible binding to streptavidin was detected, see Fig. 2A. Then, scFv genes from the third panning rounds were subcloned into the bacterial expression vector. A total of 184 monoclonal colonies from each of the two third-round selections were picked, and the soluble scFvs were assessed for binding to α-cobratoxin, revealing 290 out of the 368 (79%) displaying a binding signal against the antigen, see Fig. 2B. Among the 290 hits, 60 clones were randomly picked for further characterization, including an ENC DELFIA and DNA sequencing, see Fig. 2C for ENC data. DNA sequencing revealed that 14 of the 60 scFvs had unique CDRL3 sequences. The 13 most promising of these scFvs, based on their ranking in the ENC assay, were converted to the IgG1 format and expressed in Expi293F™ cells. A direct DELFIA assay was used to confirm that all 13 antibodies retained binding in the IgG1 format. An ENC DELFIA was employed to rank the binding of the IgGs, and along with their expression yields, 2552_02_B02 was picked as the top candidate, see Fig. 2D.

**Fig. 2.**
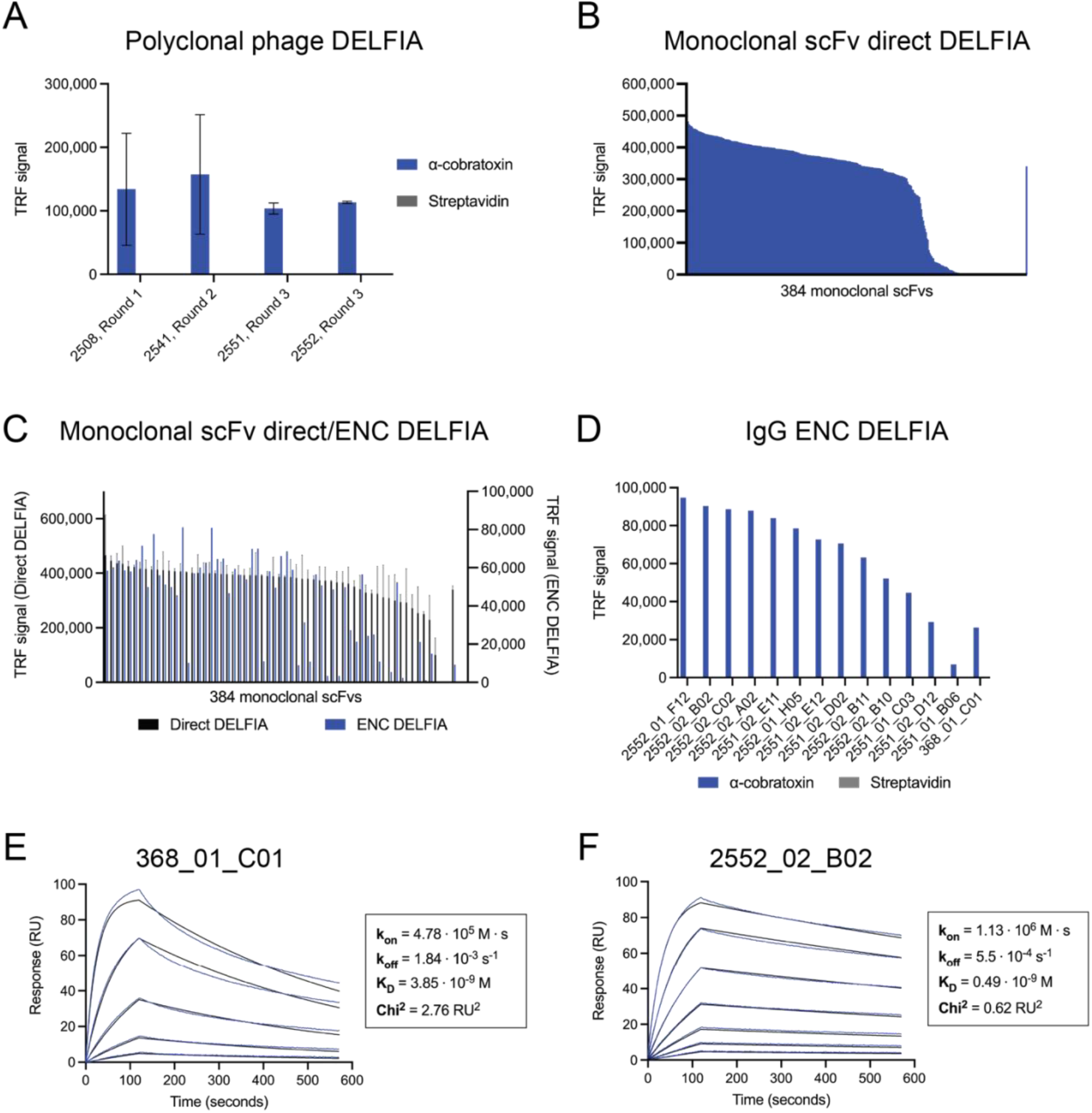
Binding, expression, and binding kinetics characterization of discovered antibodies. (**A**) Polyclonal phage DELFIA of selections outputs from three selection rounds (where antigen concentration was lowered 10-50-fold between each round) showing binding to α-cobratoxin and no binding to streptavidin. (**B**) Direct monoclonal DELFIA of 384 monoclonal scFv-containing supernatants. (**C**) Direct monoclonal DELFIA signals (in black on the left Y-axis) and ENC DELFIA signals (in blue on the right Y-axis) of the 60 monoclonal scFvs selected for sequencing. (**D**) ENC DELFIA signals of IgG-containing supernatants of the 13 clones that were converted to the IgG format. 2552_02_B02 was selected for further characterization. For (B), (C), and (D), data from the parent scFv (368_01_C01) is shown furthest to the right. (**E**) 1:1 binding model (black lines) fitted to the SPR data kinetics of parental antibody (368_01_C01) in the Fab format. (**F**) 1:1 binding model (black lines) fitted to the SPR data of the affinity matured clone (2552_02_B02) in the Fab format.

To assess the improvement in affinity derived from the exchange of the parental antibody light chain, Surface Plasmon Resonance (SPR) was employed. Here, the parent (368_01_C01) and 2552_02_B02 were reformatted as monovalent Fabs and used for the affinity measurements to avoid avidity effects. For the parent clone, the affinity (equilibrium dissociation constant, K_D_) was determined to 3.85 nM, whereas the matured clone displayed an affinity of 490 pM, representing an approximate 8-fold increase in affinity. This increase in affinity resulted from the improvement of both the on and off-rate of the original antibody, 2.4-fold and 3.3-fold, respectively (Fig. 2 E and F).

### Affinity matured IgGs show potent blocking of toxin:receptor binding interaction

To assess if the increase in affinity achieved from affinity maturation translated to improved blocking of the binding interaction between the AChR and α-cobratoxin, a receptor blocking assay was performed. A schematic representation of the assay can be seen in Fig. 3 A.

**Fig. 3.**
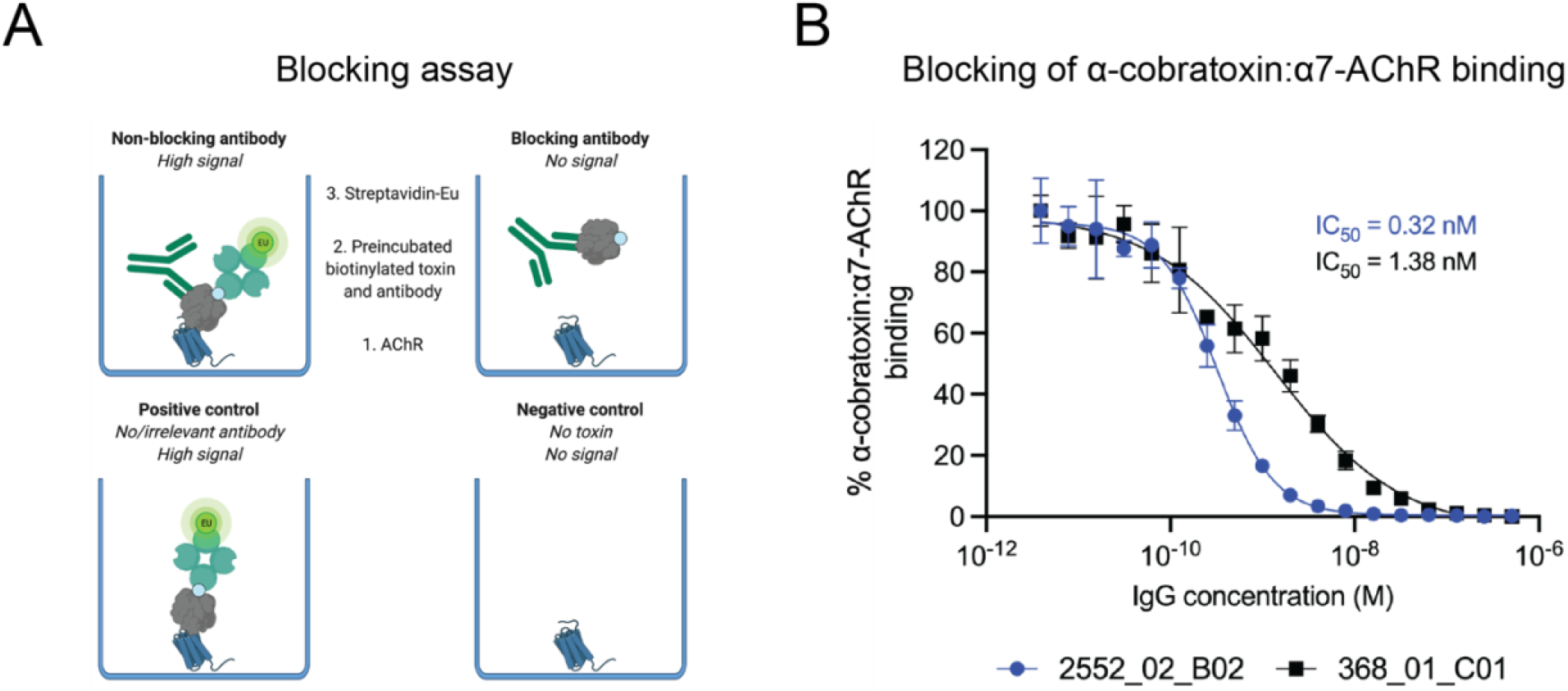
Inhibition of the binding interaction between the α7-AChR and α-cobratoxin using IgGs. (**A**) Schematic representation of a receptor blocking assay, wherein the ability of IgG antibodies to inhibit the interaction between the α7 subunit of AChR (α7-AChR) and α-cobratoxin can be quantified. (**B**) Antibodies block α-cobratoxin binding to its receptor (α7-AChR) in a concentration-dependent manner. As a negative control an IgG specific to dendrotoxins was used and showed no blocking.

Results revealed that both IgGs were able to fully abrogate the binding between the receptor subunit and α-cobratoxin, though, the affinity-matured clone was able to prevent the binding at lower concentrations than the parent antibody. The IC_50_ values were 0.32 nM (CI_95_ 0.28-0.36 nM) and 1.38 nM (CI_95_ 1.02-1.86 nM) for 2552_02_B02 and 368_01_C01, respectively (Fig. 3 B). The 8.1-fold increase in affinity to α-cobratoxin therefore resulted in a 4.3-fold improvement of IC_50_ in this blocking assay.

### Increased neutralization potency in *in vitro* neutralization assay

To assess if the increase in affinity and ability to block the α7-AChR:α-cobratoxin interaction for the affinity matured antibody also resulted in an improved ability of the antibody to protect nAChR function, functional neutralization assays were conducted using automated patch-clamp electrophysiology. First, the EC_80_ value for ACh was established (70 µM, Fig. 4 A and B), and the IC_80_ for α-cobratoxin was determined (4 nM, Fig. 4 C and D). Then, titrated 368_01_C01 and 2552_02_B02 were preincubated with α-cobratoxin and tested to examine the ability of the antibodies to neutralize the current-inhibiting activity of α-cobratoxin. As a negative control, a dendrotoxin-binding IgG was included. This irrelevant IgG showed no effects, while the α-cobratoxin-recognizing antibodies were able to fully abrogate α-cobratoxin activity. Furthermore, the affinity-matured clone, 2552_02_B02, was a more potent neutralizer with an EC_50_ of 2.6 nM (CI_95_ 2.3-2.9 nM), while the parental clone (368_01_C01) exhibited an EC_50_ of 8.1 nM (CI_95_ 6.6-10.0 nM). Relative to the concentration of α-cobratoxin used, these data indicate that 0.65 IgG molecules were needed per toxin molecule for 50% neutralization for 2552_02_B02, whereas 2.03 IgG molecules were needed per toxin to achieve the same effect with 368_01_C01. Hence, the increase in affinity between the 2552_02_B02 clone and α-cobratoxin resulted in increased functional neutralization *in vitro*.

**Fig. 4.**
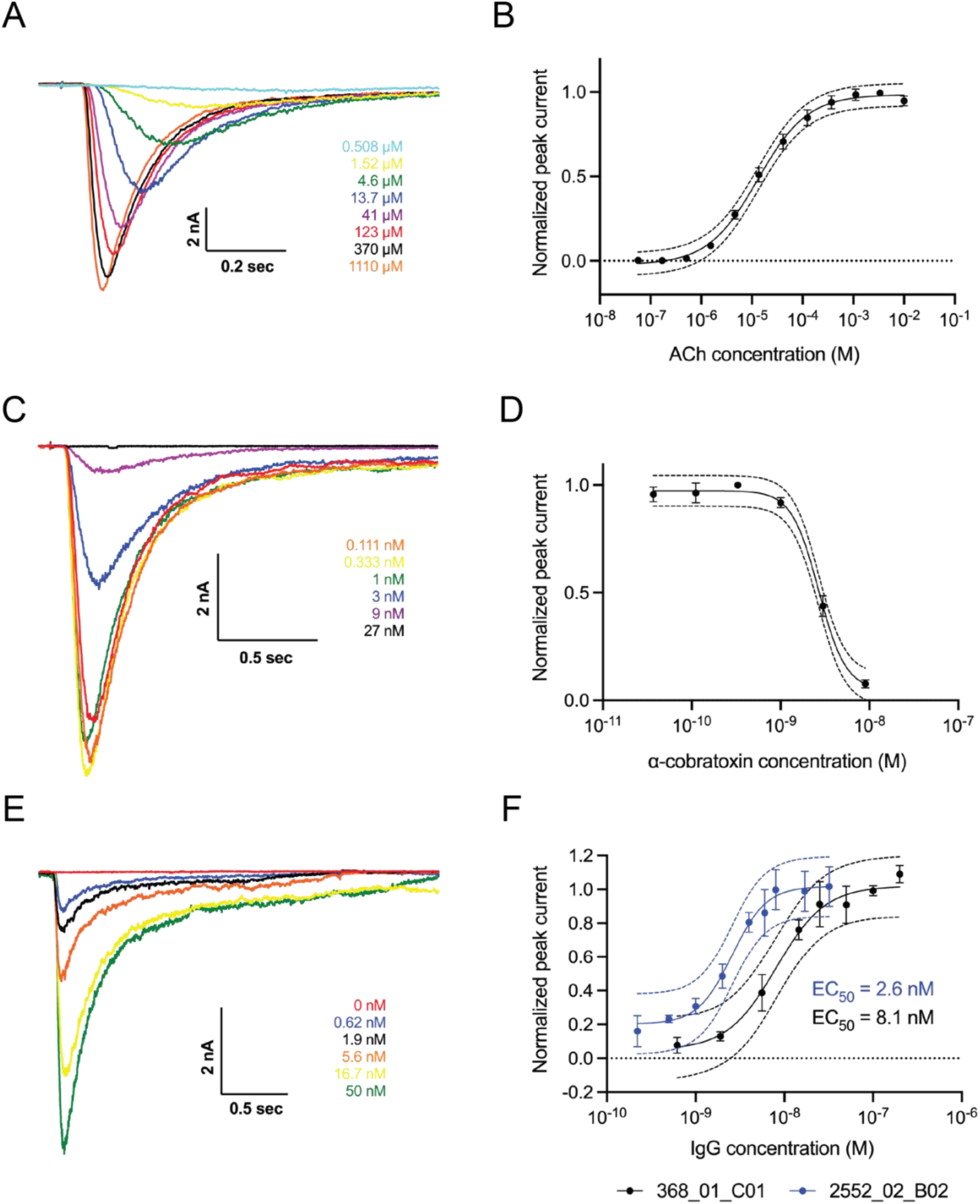
*In vitro* neutralization of inhibition of nAChR by α-cobratoxin. Automated patch-clamp experiments conducted using a QPatch (Sophion Bioscience). (**A**) Sweep plot and (**B**) concentration-response curve showing the relationship between increased ACh concentration and the measured current running across the cell membrane. 70 µM ACh was used throughout the rest of the experiments. (**C**) Sweep plot and (**D**) concentration-response curve showing how increasing concentrations of α-cobratoxin result in a decrease in the current measured. 4 nM α-cobratoxin was used, resulting in approximately 80% inhibition of the current. (**E**) Sweep plot (368_01_C01) and (**F**) concentration-response curves showing how increasing concentrations of the two IgGs preincubated with α-cobratoxin result in better protection of the nAChR, as the loss of current mediated by α-cobratoxin is prevented in a dose-dependent manner. An irrelevant IgG was used as a control, which did not prevent the inhibitory effects of α-cobratoxin.

### Affinity matured antibody neutralizes *Naja kaouthia* whole venom *in vivo*

To determine the ability of 2552_02_B02 to neutralize the effects of *N. kaouthia* whole venom, a mouse lethality assay was set up, as this is the gold standard of the WHO for the assessment of the preclinical efficacy of antivenoms. 2 LD_50_s of *N. kaouthia* whole venom were preincubated with the IgG in different molar ratios for 30 minutes before injecting the mixture i.v. in mice. As controls, venom was injected alone, in combination with a negative (irrelevant) control IgG (Nivolumab), and in combination with a commercial polyclonal antivenom as a positive control. Survival was recorded and is illustrated in Fig. 5. Mice injected with either venom alone or in combination with Nivolumab all died within 30 minutes after injection with signs of neuromuscular paralysis (Fig. 5 A). At 48 hours post injection, mice receiving venom and polyclonal antivenom were alive and showed no signs of neurotoxicity, at which point the study on this group was concluded. For the groups of 1:1 and 1:2 toxin:IgG dosages, the surviving mice at 24 hours were showing clear signs of toxicity, and it was decided to extend the observation period until 48 hours owing to the novel nature of these antibodies and the need to know whether the neutralization observed in the first hours could be reverted. At the 48-hour time point, all mice in the 1:1 ratio were dead, and only one mouse in the 1:2 toxin:IgG dosage group remained alive. Therefore, a new experiment increasing the dosage to 1:4 toxin:IgG was set up. For this group, all mice survived the 24-hour period, one mouse was dead at 48 hours, and by 72 hours post injection no further deaths had occurred (Fig 5 B). These results clearly demonstrate that the therapeutic effect of 2552_02_B02 on mice injected with a lethal amount of *N. kaouthia* whole venom were dose-dependent *in vivo*, as expected for specific antibody therapeutics. Even further, at an α-neurotoxin to IgG molar ratio of 1:4, all mice survived the observation period recommended by the WHO for this type of study (24 hours), and even at the 72-hour mark, three out of four mice had survived.

**Fig. 5.**
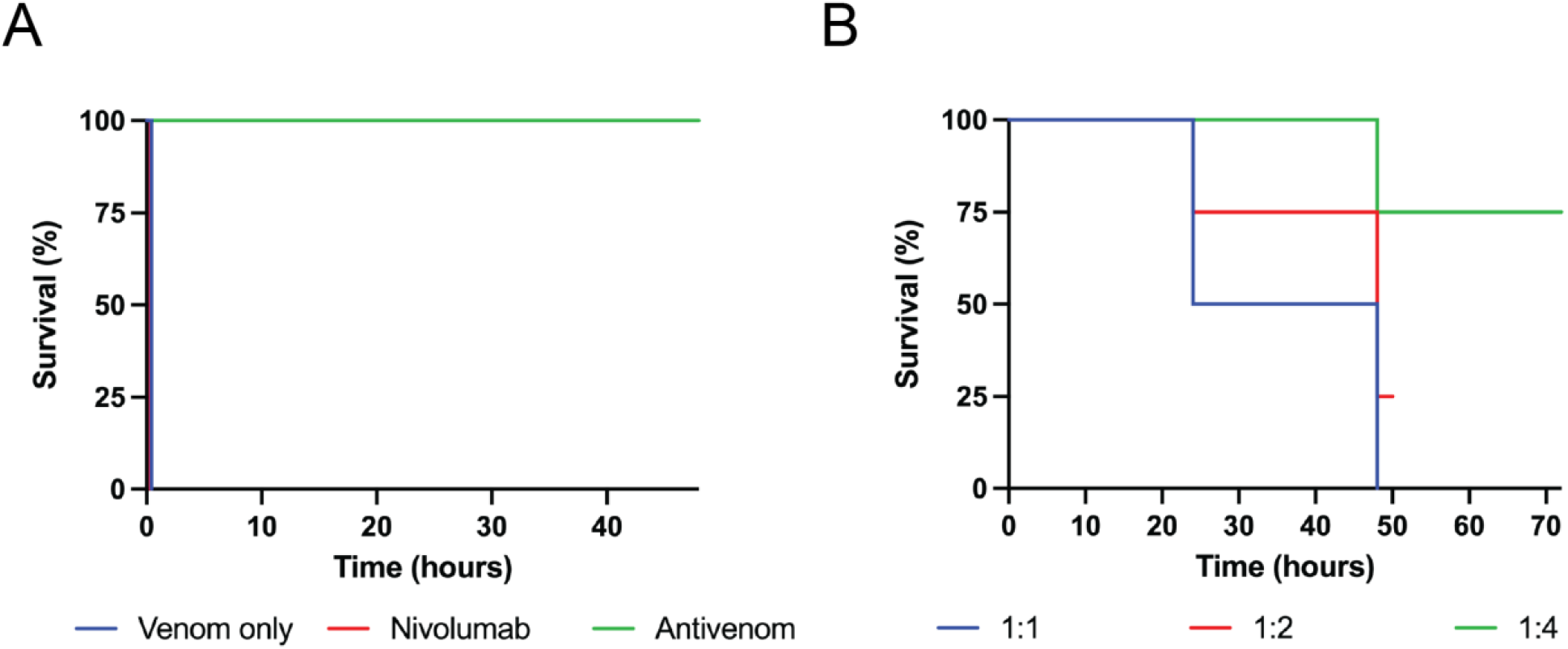
Kaplan-Meier curves showing survival of mice co-administered with *N. kaouthia* whole venom and IgG 2552_02_B02. (**A**) Illustrates that the death of the venom only and Nivolumab control groups occurred within 30 minutes after injection, whereas those receiving venom and antivenom survived the 48 hour observation period. (**B**) Illustrates the survival of the groups of mice receiving IgG and venom. 2552_02_B02 in 1:1 and 1:2 toxin:IgG molar ratios were observed for 48 hours, whereas mice receiving the 1:4 molar ratio were observed for 72 hours.

## DISCUSSION

Previously, we described the discovery of an oligoclonal mixture of human antibodies capable of neutralizing dendrotoxin-mediated neurotoxicity of black mamba venom in a rodent model *(19)*. Although the cocktail of antibodies tested in that study did neutralize whole venom, the model, using intracerebroventricular injection (i.c.v.), did not account for the effects elicited by α-neurotoxins, since their main target is the nAChR in the neuromuscular junctions. Thus the i.c.v. model is not nearly as clinically relevant as i.v. injection, which is recommended by WHO as the standard for assessing antivenoms. Another study has reported *in vivo* neutralization of α-cobratoxin-induced lethality by a V_H_H and a V_H_H2-Fc following intraperitoneal injection in mice *(26)*. However, to date no study has successfully demonstrated the neutralization of lethality caused by a whole venom (or a purified toxin) preincubated with a recombinant monoclonal IgG antibody following i.v. injection.

In this study, we demonstrate that a recombinant human monoclonal IgG antibody, discovered and optimized entirely *in vitro* by phage display technology, was able to neutralize lethality in mice challenged i.v. with whole venom from *N. kaouthia*. This clearly showcases the utility of *in vitro* selection methods for the discovery of efficacious antivenom antibodies against animal toxins with reduced immunogenicity, which may be challenging for traditional *in vivo* based discovery approaches *(27)*. Moreover, this study has also elucidated the mechanism of action of the neutralizing antibody using receptor blocking and automated patch clamp electrophysiology assays, which revealed that the antibody could abrogate neurotoxicity by preventing the medically most important toxin, α-cobratoxin, from interacting with the nAChR. This first report of a recombinant monoclonal antibody neutralizing whole venom from a snake following i.v. injection thus presents an integrated approach for future discovery and evaluation of recombinant antibodies against toxins from snake and other animal venoms. In this relation, it is, however, important to note that the neutralization of this particular venom by a single IgG cannot be extrapolated to all other snake venoms due to their complex composition of different medically relevant toxin families that each may require one or more antibodies for neutralization *(28)*. In many cases, co-administration with other antibodies or small molecule inhibitors, such as varespladib, batimastat, or marimastat *(29, 30)*, might be necessary to achieve full protection *(18)*. Furthermore, the biophysical and pharmacokinetic properties of the IgG reported in this study have not been investigated and therefore these properties remain unknown. Antibody pharmacokinetics play a significant role in drug efficacy *(31)*, while antibody biophysics can have major influence on antibody manufacturability and stability *(32–34)*. Additional investigation of these properties is therefore warranted prior to further preclinical and clinical assessments.

Finally, a regulatory uncertainty currently exists regarding whether recombinant antivenoms based on monoclonal or oligoclonal antibodies will be regulated as blood products, similar to existing plasma-derived antivenoms, or whether recombinant antivenoms will be viewed as biotherapeutic products to be regulated as biopharmaceuticals by relevant authorities.

Establishment of a regulatory framework for recombinant antivenom products is thus a necessity for bringing such new snakebite envenoming therapies swiftly to the clinic.

Nonetheless, the advances in the discovery, optimization, and assessment of monoclonal antibodies against snake toxins described in this study represent an important technical milestone towards the application of *in vitro* developed recombinant antivenoms as a therapeutic intervention in snakebite envenoming in the future.

## MATERIALS AND METHODS

### Study design

The original objective of the study was to discover human monoclonal IgGs that could neutralize α-cobratoxin *in vivo*. When proven unsuccessful, the aim of the study became to test if affinity maturation using chain-shuffling of an α-cobratoxin-specific IgG antibody would result in improved *in vitro* and *in vivo* neutralization and to test if administration of only α-cobratoxin specific IgGs would enable full neutralization of *N. kaouthia* whole venom *in vivo*. The initial discovery and subsequent affinity maturation of α-cobratoxin specific antibodies was carried out *in vitro*. The following assessment of binding was conducted on the antibodies in different formats (scFv, Fab, and IgG) using several different assays (direct DELFIA, ENC DELFIA, and SPR). The following assessment of *in vitro* neutralization was conducted using two different methods (automated patch-clamp and receptor blocking) at different research institutions, both reaching similar conclusions that the IgGs were neutralizing. For the *in vivo* neutralization of lethality assays, mice were randomized into treatment groups of 3-4 mice per group (group size determined based on previous studies) with predefined endpoint at 24 hours; in some cases, owing to the novel nature of the antibodies, the endpoint of these *in vivo* assays was extended.

All assays were conducted with 2-5 repeats (except for animal experiments) in at least technical triplicates with negative controls included in all experiments and positive controls included when possible.

### Toxin preparation

α-cobratoxin was obtained in lyophilized form from Latoxan SAS, France. The toxin was reconstituted in phosphate buffered saline (PBS) and biotinylated using a 1:1 (toxin:biotinylation reagent) molar ratio as previously described *(19)*. Following biotinylation, Amicon® Ultra-4 Centrifugal Filter Units with a 3 kDa membrane were used for purification of the biotinylated toxin. Purification was performed at 8 °C and consisted of three washes of 4 mL PBS. The protein concentration was measured by the absorbance at 280 nm using a Nanodrop and adjusted using the extinction coefficient. The degree of biotinylation was analyzed using MALDI-TOF in a Proteomics Analyzer 4800 Plus mass spectrometer (Applied Biosystems).

### Initial discovery and assessment of parent clone

The initial discovery of the parent antibody clone 368_01_C01 was performed by panning the IONTAS phage display library (diversity of 4 × 10^10^ human scFv clones) against biotinylated α-cobratoxin captured by streptavidin in a Maxisorp vial, followed by subcloning and expression of scFv genes in BL21(DE3) *E. coli* and DELFIA-based screening of the scFv-containing supernatants as previously described *(19)*. Thirty-eight binding clones were cherry-picked and sequenced (Eurofin Genomics sequencing service) using the S10b primer (GGCTTTGTTAGCAGCCGGATCTCA). The binding strengths of the clones were ranked using an expression-normalized capture (ENC) assay, and the top six scFvs displaying the highest binding signals were reformatted into the IgG format and expressed in Expi293™ cells (Thermo Fisher) and subsequently purified using an Äkta Pure system (GE Healthcare) as previously described *(19)*. The functionality of purified IgGs was confirmed using a DELFIA-based binding assay *(19)*.

### Library generation using chain-shuffling

The V_H_ region of antibody 368_01_C01 was PCR amplified from pINT3 plasmid DNA using pINT3 Nco FWD (TCTCTCCACAGGCGCCATGG) and IgG1 CH1 Xho Rev (CCCTTGGTGGAGGCACTCGAG) primers using Platinum™ SuperFi II Green PCR Master Mix (Invitrogen, 12369010). The PCR product was cloned into pIONTAS1 *(23)* vector harboring the naïve V_L_ lambda and kappa chain libraries using *NcoI* and *XhoI* restriction endonucleases. Ligation reactions were carried out for 16 hours at 16 °C and contained 160 ng of insert and 400 ng of vector DNA in a total volume of 40 µL. Ligations were purified using the MinElute PCR Purification Kit (Qiagen, 28004), and eluted in 10 µL nuclease free water. The purified ligation product was transformed into 200 µL of electrocompetent TG1 cells (Lucigen, 60000-PQ763-F) followed by addition of 6 mL of recovery medium (Lucigen, F98226-1) and incubation at 37 °C for one hour at 280 rpm rotation. Cells were plated on 2xTY agar plates supplemented with 100 µg/mL ampicillin and 2% glucose. Dilutions of the transformations were also plated to determine library size, which was 1.01 × 10^8^ for the lambda library and 1.67 × 10^8^ for the kappa library with more than 96% of the transformants being positive for insertion of heavy chain insert, as determined by colony PCR.

### Library rescue and solution-based phage display selection

Rescue of phages from the chain-shuffled libraries and the three rounds of selections were performed as described elsewhere *(23)*, except that the phages were not concentrated using PEG precipitation, but phage-containing supernatants were used directly for selections. Deselection of streptavidin-specific phages was performed before each round of selection using 80 µL of streptavidin-coated Dynabeads (Invitrogen, M-280). Additionally, the selections were conducted using biotinylated α-cobratoxin that was captured using 80 µL of streptavidin-coated Dynabeads (Invitrogen, M-280). The concentration of α-cobratoxin was decreased through the three rounds of selections starting at 10 nM in the first round and ending at 20 pM in the third round. The kappa and lambda libraries were mixed before the first round of selections.

### Subcloning, primary screening, and sequencing of scFvs

Subcloning of the α-cobratoxin binding selection output into pSANG10-3F and primary screening of candidates was performed as described elsewhere *(19)*. In brief, scFv genes from the selection outputs were subcloned from the pIONTAS1 phagemid vector to the pSANG10-3F expression vector using *NcoI* and *NotI* restriction endonuclease sites and transformed into *E. coli* strain BL21 (DE3) (New England Biolabs). From each of the two subcloned selection outputs, 184 colonies were picked and expressed in 96 well plates. The scFvs were assessed for their binding to biotinylated α-cobratoxin (5 µg/mL) indirectly immobilized on black Maxisorp plates (Nunc) with streptavidin (10 µg/mL) using a DELFIA-based assay. In total, 60 clones binding to α-cobratoxin were cherry-picked and sequenced (Eurofin Genomics sequencing service) using S10b primer (GGCTTTGTTAGCAGCCGGATCTCA). The antibody framework and CDR regions were annotated, and light chain CDR3 regions were used to identify 14 unique clones.

### IgG expression and purification

V_H_ and V_L_ genes of 13 unique α-cobratoxin-binding scFvs were converted to the IgG1 format as previously described *(19)*. The binding of the IgGs was confirmed and ranked using and expression-normalized capture (ENC) assay. Briefly, black Maxisorp plates (Nunc) were coated overnight with an anti-human IgG (Jackson ImmunoResearch, 109-005-098). Plates were washed thrice with PBS and blocked with PBS supplemented with 3% milk protein. Plates were washed thrice with PBS and 0.25x unpurified IgG-containing culture supernatant in PBS supplemented with 3% milk protein was added before incubating for one hour at room temperature. Plates were washed thrice with PBS-T and thrice with PBS before adding either 1nM or 100 pM biotinylated α-cobratoxin in PBS supplemented with 3% milk protein to each well. After one hour of incubation, the plates were washed thrice with PBS-T and thrice with PBS. Then, 1 µg/mL of Europium-labeled Streptavidin (Perkin Elmer, 1244-360) in DELFIA Assay Buffer (Perkin Elmer, 4002-0010) was added. Following 30 minutes of incubation, plates were washed thrice with PBS-T and thrice with PBS, and DELFIA Enhancement Solution (Perkin Elmer, 4001-0010) was added for detection of binding. Based on these results, the two top clone, 2552_02_B02, was expressed and purified as described previously *(19)*.

### Fab expression and purification

V_H_ and V_L_ genes of 13 unique α-cobratoxin-binding scFvs were converted to the Fab format as performed for IgGs as described previously *(19)*, except the Fab-vector, pINT12, was used instead of the pINT3 IgG1 vector.

### Surface plasmon resonance

The binding affinity of the discovered antibodies for α-cobratoxin was determined using Surface Plasmon Resonance (SPR) (BIAcore T100, GE Healthcare). All measurements were performed at 25 °C using 10 mM HEPES, 150 mM NaCl, and 3 mM EDTA at pH 7.4 as running buffer.

Immobilisation of α-cobratoxin on CM5 sensor chips (Cytiva, BR100530) was performed by amine coupling using 1-Ethyl-3-(3-dimethylaminopropyl)carbodiimide (EDC)/ N-hydroxysuccinimide (NHS) surface activation followed by injection of 5 µg/mL of α-cobratoxin in 10 mM NaOAc pH 4 to btain a final immobilization level of 23 response units (RU). The sensor chip was inactivated using ethanolamine. The Fabs in concentrations ranging from 81 nM to 390 pM were injected at 40 µL/minute for 120 seconds and dissociation was recorded for 450 seconds. Following dissociation, the sensor was regenerated using two injections (15-20 seconds) of 20 mM NaOH. Measurements were conducted using 5-7 analyte concentrations for each antibody. The blank subtracted data was analyzed using the BIAcore T100 Evaluation Software employing a 1:1 Langmuir binding model.

### Receptor blocking DELFIA

The receptor blocking assay was adapted from Ratanabanangkoon *et al. (35)*. Black Maxisorp plates (Nunc) were coated overnight with 100 µL of 5 µg/mL human α7-acetycholine receptor chimera (adapted from *(36)*) in PBS. Plates were washed thrice with PBS and blocked with 1% BSA in PBS. 4 nM of biotinylated α-cobratoxin with various concentrations of 368_01_C01, 2552_02_B02, or a negative control IgG specific to dendrotoxins in 0.1% BSA was prepared and preincubated for 30 minutes at room temperature. Plates were washed thrice, and 100 µL of the preincubated toxin and antibody mixture was added to the blocked wells. Following incubation for one hour, the plates were washed thrice with PBS-T (PBS, 0.1% Tween-20) and thrice with PBS, and 100 µL of 1 µg/mL of Europium-labeled Streptavidin (Perkin Elmer, 1244-360) in 0.1% BSA was added. Following 30 minutes of incubation, plates were washed thrice with PBS-T and thrice with PBS and 100 µL of DELFIA Enhancement Solution (Perkin Elmer, 4001-0010) was added to each well. Signals were measured using a VICTOR Nivo Multimode Microplate Reader using excitation at 320 nm and emission at 615 nm. Each antibody concentration was tested in quadruplicate.

### Electrophysiology

Planar whole-cell patch-clamp experiments were carried out on a QPatch II automated electrophysiology platform (Sophion Bioscience), where 48-channel patch chips with 10 parallel patch holes per channel (patch hole diameter ∼1 μm, resistance 2.00 ± 0.02 MΩ) were used.

The cell line used was a human-derived Rhabdomyosarcoma RD cell line (CCL-136, from ATCC), endogenously expressing the muscle-type nicotinic acetylcholine receptors (nAChR), composed of the α1, β1, δ, γ, and ε subunits. The cells were cultured according to the manufacturer’s guideline, and on the day of the experiment, enzymatically detached from the culture flask and brought into suspension.

For patching, the extracellular solution contained: 145 mM NaCl, 10 mM HEPES, 4 mM KCl, 1 mM MgCl_2_, 2 mM CaCl_2_, and 10 mM glucose, pH adjusted to 7.4 and osmolality adjusted to 296 mOsm. The intracellular solution contained: 140 mM CsF, 10 mM HEPES, 10 mM NaCl, 10 mM EGTA, pH adjusted to 7.3, and osmolality adjusted to 290 mOsm.

In the experiments, an nAChR-mediated current was elicited by 70 µM acetylcholine (ACh, Sigma-Aldrich), approximately the EC_80_ value, and after compound wash-out, 2 U acetylcholinesterase (Sigma-Aldrich) was added to ensure complete ACh removal. The ACh response was allowed to stabilize over three ACh additions, before the fourth addition was used to evaluate the effect of α-cobratoxin (4 nM α-cobratoxin, reducing the ACh response by 80%), preincubated with varying concentrations of IgGs. α-cobratoxin and IgGs were preincubated at room temperature for at least 30 minutes before application, and the patched cells were preincubated with α-cobratoxin and IgG for 5 minutes prior to the fourth ACh addition. As a negative control, an IgG specific to dendrotoxins was included.

The inhibitory effect of α-cobratoxin was normalized to the full ACh response (fourth response normalized to third response), plotted in a non-cumulative concentration-response plot, and a Hill fit was used to obtain EC_50_ values for each IgG. The data analysis was performed in Sophion Analyzer (Sophion Bioscience) and GraphPad Prism (GraphPad Software).

### Animals

*In vivo* assays were conducted in CD-1 mice of both sexes of 18-20 g body weight, supplied by Instituto Clodomiro Picado, following protocols approved by the Institutional Committee for the Use and Care of Animals (CICUA), University of Costa Rica (approval number CICUA 82-08). Mice were housed in cages in groups of 3-4 and were provided food and water *ad libitum*.

### *In vivo* preincubation experiments

The neutralization activity of the naïve IgGs against α-cobratoxin was tested by i.v. injection in groups of three mice. 4 µg of α-cobratoxin (corresponding to 2 LD_50_s) and 150 µg of corresponding IgG (α-cobratoxin:IgG = 1:2 molar ratio) were dissolved in PBS, preincubated (30 min at 37 °C), and injected in the caudal vein, using an injection volume of 100 µL. Control mice were injected with either anti-lysozyme IgG and α-cobratoxin or α-cobratoxin alone. Deaths were recorded and Kaplan-Meier curves were used to represent mouse survival along time.

For the affinity matured clone, similar *in vivo* experiments were conducted, except the IgG was preincubated with 9.12 µg of *N. kaouthia* whole venom (corresponding to 2 LD_50_s) at a 1:1, 1:2, and 1:4 α-neurotoxin:IgG molar ratio. For calculating molar ratios, it was estimated that 55% of *N. kaouthia* venom consist of α-neurotoxins, based on a toxicovenomic study of the venom *(7)*. All injections were performed as described above on groups of four mice. Control mice were injected with either Nivolumab (irrelevant IgG control) incubated with *N. kaouthia* venom or *N. kaouthia* venom alone. As a positive control for *N. kaouthia* venom neutralization, Snake Venom Antiserum from VINS Bioproducts Limited (Batch number: 01AS13100) was used. According to the manufacturer, the potency of this antivenom against the venom of *Naja naja* is 0.6 mg venom neutralized per mL antivenom. Since no information is provided on the neutralization of *N. kaouthia* venom, we used a ratio of 0.2 mg venom per mL antivenom to ensure neutralization. Survival was monitored for 24-72 hours, and results are presented in Kaplan-Meier curves.

## Acknowledgments

LL is thankful to Georgia Bullen for general guidance in the laboratory as well as to Yessica Wouters for help with data analysis. Birte Svensson is thanked for discussions on SPR analysis. Figure 3 A was created using BioRender.com.

## Funding

Villum Foundation grant 00025302 (AHL)

The European Research Council (ERC) under the European Union’s Horizon 2020 research and innovation programme grant no. 850974 (AHL)

The Novo Nordisk Foundation (NNF16OC0019248) (AHL)

The Hørslev Foundation (203866) (AHL)

Olsens Mindefold (LL)

Marie og M.B. Richters Fond (LL)

Niels Bohr Fondet (LL)

Torben og Alice Fritmodts Fond (LL))

William Demant Fonden (LL)

Otto Mønsteds Fond (LL).

Knud Højgaards Fond (LL)

Rudolph Als Fondet (LL)

Augustinus Fonden (LL)

Tranes Fond (LL)

## Author contributions

Conceptualization: AHL, LL, AKV

Methodology: AHL, LL, AKV, JMC, PS, KB, JMG, BL

Investigation: AHL, LL, UP, JW, PV, KB, EWM, ASA, SO, DTG, AML, ML, RAL, HA, JMC, JMG, BL, MSM

Visualization: LL, KB

Writing – original draft: LL, AHL, KB

Writing – review & editing: LL, AHL, AKV, JMG, BL, PS

